# Dual fates of exogenous tau seeds: lysosomal clearance vs. cytoplasmic amplification

**DOI:** 10.1101/2022.01.03.474802

**Authors:** Sourav Kolay, Anthony R. Vega, Dana A. Dodd, Valerie A. Perez, Omar M. Kashmer, Charles L. White, Marc I. Diamond

**Author notes:** Corresponding Author: Marc Diamond, M.D., NL10.120, 6000 Harry Hines Blvd. Dallas, TX. 75390.

## Abstract

Tau assembly propagation from the extracellular to intracellular space of a cell may underlie neurodegenerative tauopathies. The first step involves tau binding to heparan sulfate proteoglycans on the cell surface, followed by macropinocytosis. Pathological tau assemblies are thought to exit the vesicular compartment as “seeds” for replication in the cytoplasm. Tau uptake is highly efficient, but only ∼1-10% of cells that take up aggregates exhibit seeding. To investigate the basis for this observation, we used fluorescently tagged full-length (FL) tau fibrils added to native U2OS cells, and “biosensor” cells expressing FL tau or repeat domain fused to mClover (Clo). FL tau-Clo bound tubulin, but seeds triggered its aggregation in multiple locations simultaneously in the cytoplasm, generally independent of visible exogenous aggregates. Most exogenous tau trafficked to the lysosome, but imaging revealed a small percentage that slowly and steadily accumulated in the cytosol. Intracellular expression of Gal3-mRuby, which binds intravesicular galactosides and forms puncta upon vesicle rupture, revealed no evidence of vesicle damage following tau exposure. In fact, most seeded cells had no evidence of lysosome rupture. However, live cell imaging indicated that cells with pre-existing Gal3-positive puncta exhibited seeding at a slightly higher rate than the general population, indicating a potential role for vesicle instability as a predisposing factor. Clearance of tau seeds occurred rapidly in both vesicular and cytosolic fractions. Bafilomycin inhibited vesicular clearance, whereas MG132 inhibited cytosolic clearance. Tau seeds that enter the cell thus have at least two fates: lysosomal clearance that degrades most tau, and entry into the cytosol, where seeds replicate, and are cleared by the proteasome.

## Introduction

Multiple lines of experimental evidence suggest that tau protein triggers neurodegeneration after intracellular accumulation in ordered assemblies. Myriad tauopathies are linked to distinct assembly structures, and include Alzheimer’s disease (AD), frontotemporal dementia, and chronic traumatic encephalopathy, among many others (1). The transcellular propagation and faithful replication of unique assembly structures, or “strains,” appears to underlie the characteristic progression patterns of specific tauopathies (2). Tau propagation presumably involves three steps: uptake of an assembly into the cell; amplification and maintenance of the aggregated state; and exit from the cell. We originally observed that tau binding to heparan sulfate proteoglycans (HSPGs) on the cell surface underlies uptake via macropinocytosis, and is mediated by specific sulfation patterns (3, 4). After uptake into macropinosomes, an assembly must make contact with endogenous tau in the cytoplasm to serve as a template and amplify a specific structure. Several reports suggest that this might happen after vesicle rupture (5) (6), but the details are unclear. Finally, when tau seeds enter the cytoplasm, it is unknown how they are degraded. In this study, we used cultured cells to dynamically visualize tau uptake and seeding, to track the relationship to vesicle processing and rupture, and to determine mechanisms of seed degradation in the vesicular vs. cytoplasmic fractions.

## Results

### Tau binds tubulin and is recruited to aggregates in the cytosol

Wild-type full-length tau undergoes seeded aggregation inefficiently in cultured cells, and thus we studied full-length (2N4R) tau containing a disease-associated P301S mutation, fused to mClover3 (FL tau-Clo) (Fig. 1A). We stably expressed FL tau-Clo in U2OS cells, where it colocalized with tubulin (Fig. 1B). We then directly imaged U2OS biosensor cells that were exposed to exogenous full-length (2N4R) tau fibrils covalently labeled with Alexa647, tracking FL tau-Clo puncta formation over time. Similar to our original observations (11), a minority of cells exhibited induced aggregation of tau, which predominated in the cytoplasm (Fig. 1C). Time-lapse imaging with an IN Cell Analyzer 6000 (GE) also revealed intracellular aggregation in the cytoplasm (Fig. 1D), and in imaging dynamic inclusion formation in ∼50 cells, we often observed aggregates forming simultaneously throughout the cell (Supplemental Movie). For nascent intracellular aggregates, we observed no significant colocalization with Alexa647-labeled exenous tau. This raised the question of how tau traffics into the cytosol to serve as a seed.

**Figure 1:**
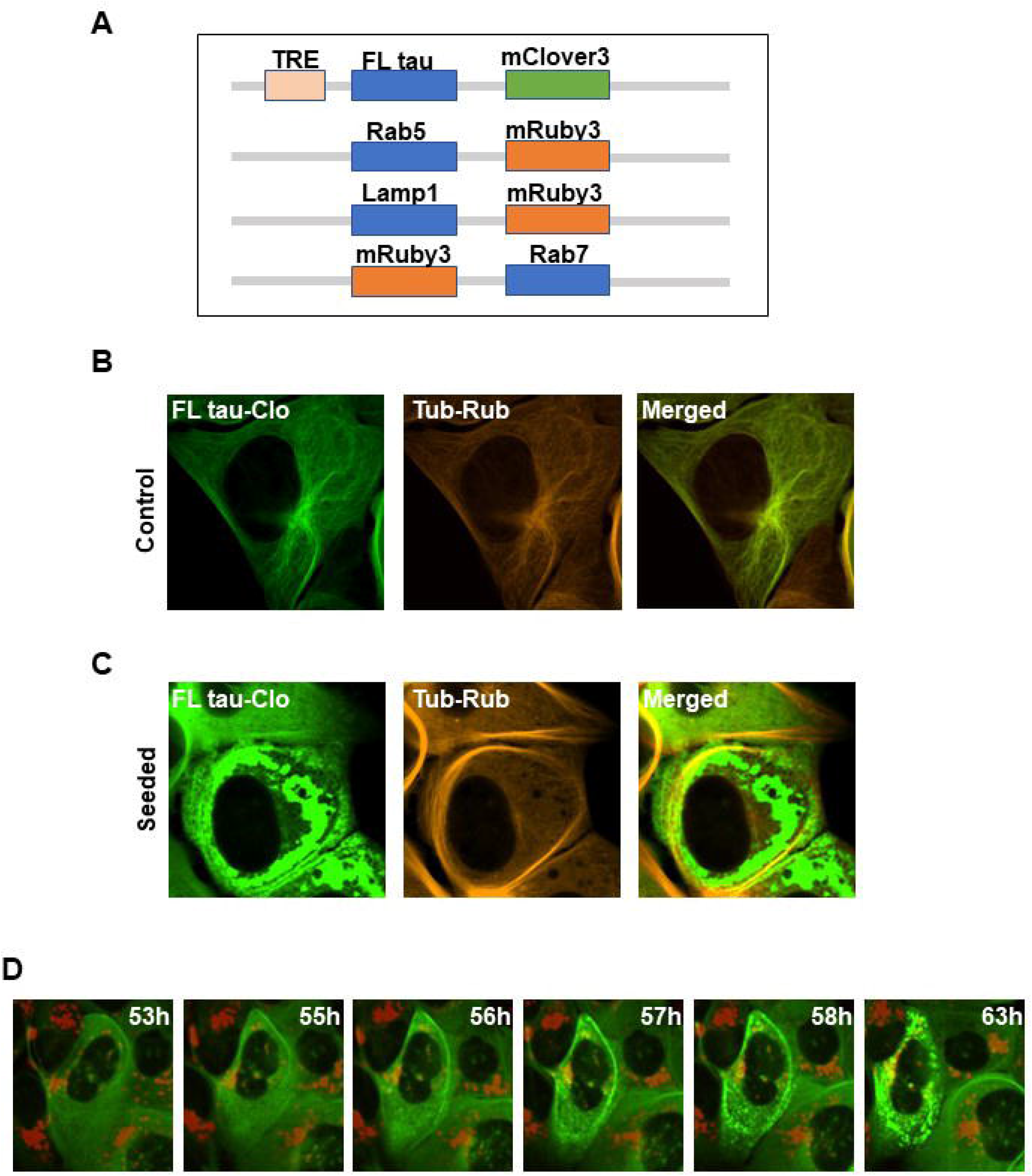
Tau aggregation initiates in the cytoplasm. (A) Diagram of constructs used for this study. (B) U2OS cells expressing FL tau-Clo containing the P301S mutation, and stained for tubulin, without exogenous fibril addition. FL tau-Clo colocalizes with tubulin. (C) With exogenous fibril addition, tau re-distributes to inclusions in the cytoplasm. (D) Recombinant FL tau fibrils labeled with AF647(red) were applied to the U2OS cells expressing FL tau-Clo (green) and imaged using a GE IN Cell Analyzer 6000 to track tau aggregation over time (labeled). Note rapid and simultaneous development of tau inclusions throughout the cytoplasm.

### Internalized tau aggregates traffic to the endolysosomal system

To study the fate of internalized tau, we exposed U2OS to FL tau fibrils labeled with Alexa647, imaging them repeatedly over 2 days with high content microscopy. We first tested for colocalization with vesicles by stably expressing mRuby3 fusions to Rab5 (to mark early endosomes, Fig. 1A, 2A), Rab7 (to mark late endosomes, Fig. 1A, 2B) and LAMP1 (to mark lysosomes, Fig. 1A, 2C). Tau progressively colocalized with these markers, especially LAMP1. We concluded that most internalized tau entered the endolysosomal pathway. When we tracked seeding in U2OS biosensor cells expressing FL tau-Clo using LAMP1, we observed no significant colocalization of emergent tau puncta with the lysosome, which was largely in a separate compartment vs. the induced FL-tau-Clo puncta (Fig. 3A). Full length tau seeding is inefficient enough to make capture of large numbers of seeding events relatively difficult. We therefore performed the same experiment using RD tau-Clo to image a high number of seeded cells (Fig.

**Figure 2:**
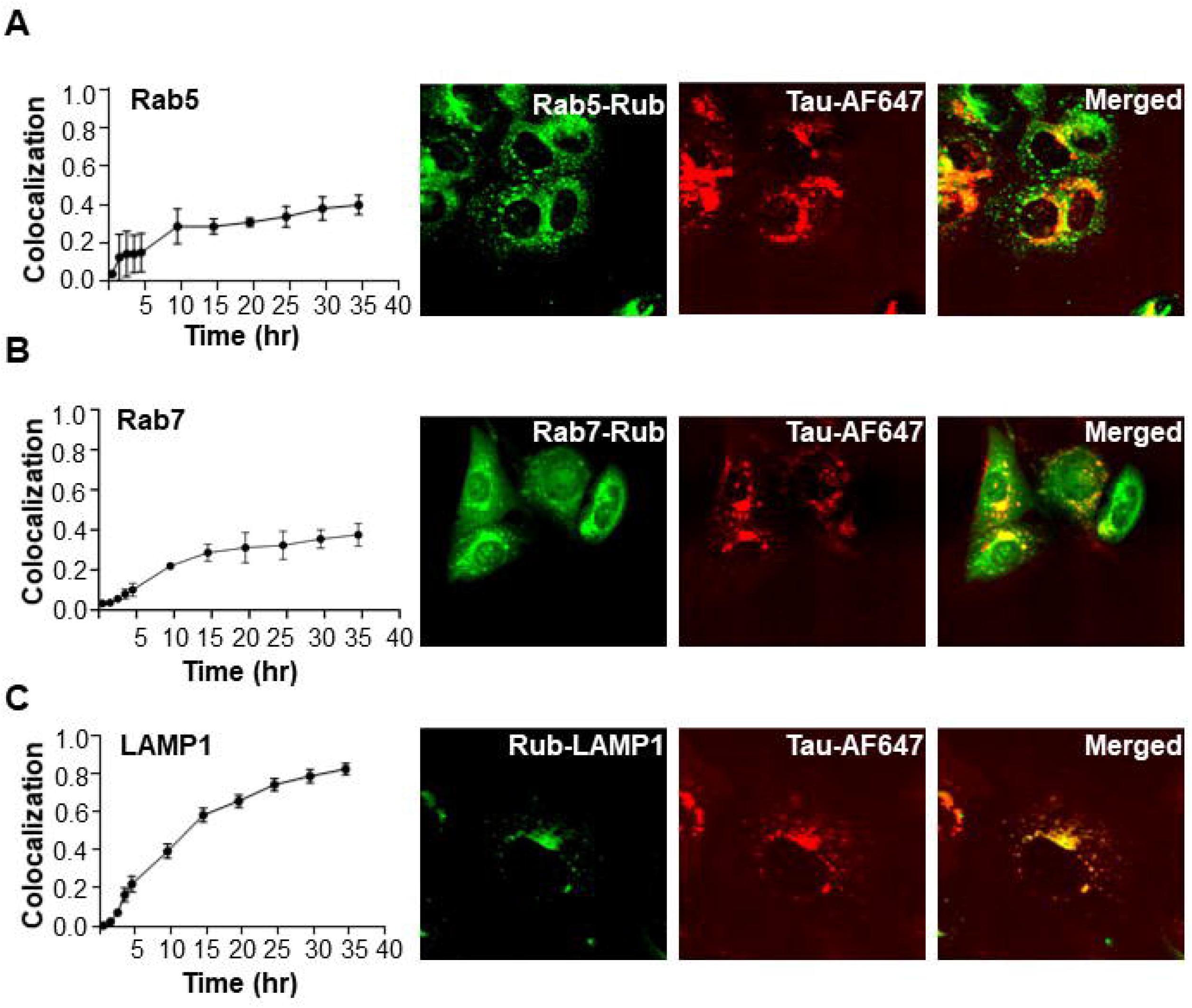
Exogenous tau fibrils co-localize with the lysosome. U2OS cells were treated with FL tau fibrils labeled with AF647. Markers of early (Rab5), late (Rab7) and lysosome (LAMP1) were expressed as mRuby (Rub) fusions to track colocalization with internalized fibrils (see also Figure 1A). Cells were tracked over time using high content microscopy. Colocalization ratio refers to the fractional colocalization of the two fluorescent signals. Representative endpoint images are shown. (A) Colocalization of tau with Rab5-Rub increases over time, but never exceeds ∼0.4 over 36h. (B) Colocalization of tau with Rub-Rab7 reaches ∼0.4 over 36h. (C) Colocalization with Lamp1-Rub reaches ∼0.8 over 36h. n>200 cells were imaged iteratively for each condition. Error bars = S.D.

**Figure 3:**
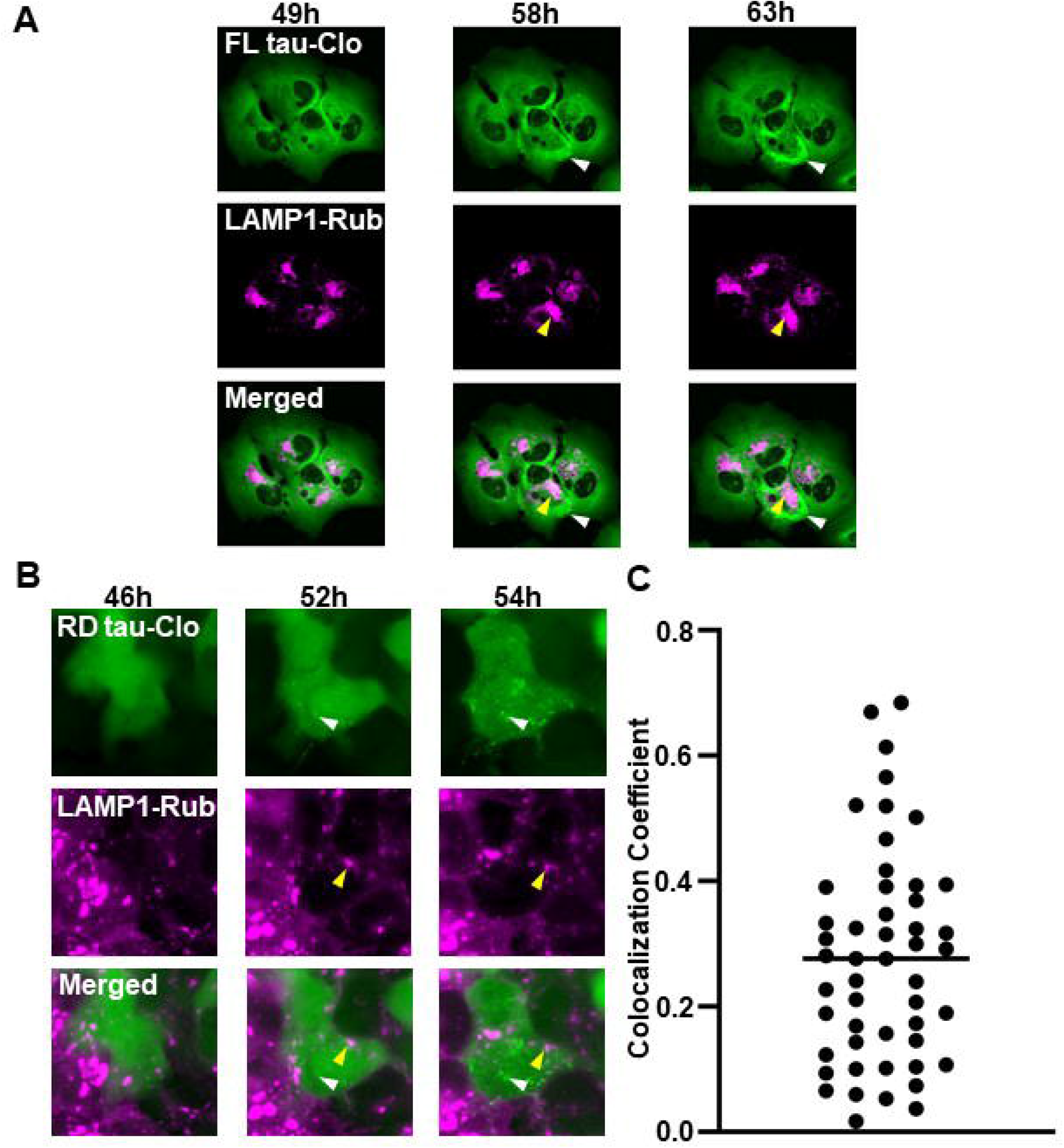
Tau inclusion formation is independent of the lysosome. (A) U2OS cells expressing FL tau-Clo and LAMP1-Rub were treated with exogenous fibrils. Cells were observed over time for coincidence of LAMP1-Rub signal and FL tau-Clo inclusion formation. A representative image shows the seeded tau (white arrowhead) and LAMP1 signal (magenta). No colocalization was observed between tau aggregates (white arrow) and lysosome (yellow arrow). Time after tau fibril addition is indicated above each column. The images are representative of 5 seeded cells studied. (B) U2OS cells expressing RD Tau-Clo and LAMP1-Rub were treated with tau fibril and followed over time for coincidence of LAMP1 signal and RD Tau-Clo aggregation. No colocalization was observed between tau aggregates (white arrowhead) and the LAMP1 marking lysosome (yellow arrowhead). The images are representative of 50 seeded cells. (C) Quantification of colocalization of LAMP1-Rub with RD tau-Clo aggregates. The graph shows colocalization coefficient (Mander’s overlap coefficient) from 50 seeded cells, and the bar indicates the mean.

3B). Similar to our observation with full length tau, we observed no significant colocalization between RD tau-Clo aggregates and LAMP1(Fig.3C). Thus, although most internalized tau wound up in the lysosome, this seemed unlikely to the be the primary location of seeding.

### Tau seeding is independent of the lysosome

Since we did not observe significant colocalization of newly formed tau puncta with vesicle markers, we hypothesized that tau seeds might be released into the cytoplasm from a vesicular pool. After exposing U2OS cells to FL tau fibrils labeled with Alexa647 we detected diffuse fluorescence in the cytoplasm at 20h (Fig. 4A). We then used live cell imaging to monitor hundreds of cells exposed to FL tau fibrils tagged with Alexa647. We observed a small but steady increase of cytosolic Alexa647 signal, whereas transferrin-Alexa647 did not increase in signal following a similar exposure protocol (Fig. 4B). To test our observations by a different approach, we used cell fractionation based on differential centrifugation to measure tau levels in cytosol vs. organelle (vesicle) fractions (Fig. 4C). We confirmed the accuracy of the fractionations using western blot against GAPDH (cytosol); VDAC (organelle); LAMP1 (organelle); and Lamin B1 (nucleus) (Fig. 4D). We attempted to quantify tau protein levels via western blot, ELISA, and mass spectrometry, but they were too low for reliable measurements. We next monitored seeding activity in the cytoplasm by transducing lysate into a well-characterized biosensor assay based on a next-generation cell line, v2L (12,13). We observed a steady increase in cytosol seeding activity over time (Fig. 4E). Tau seeds thus steadily move from vesicles to the cytosol.

**Figure 4:**
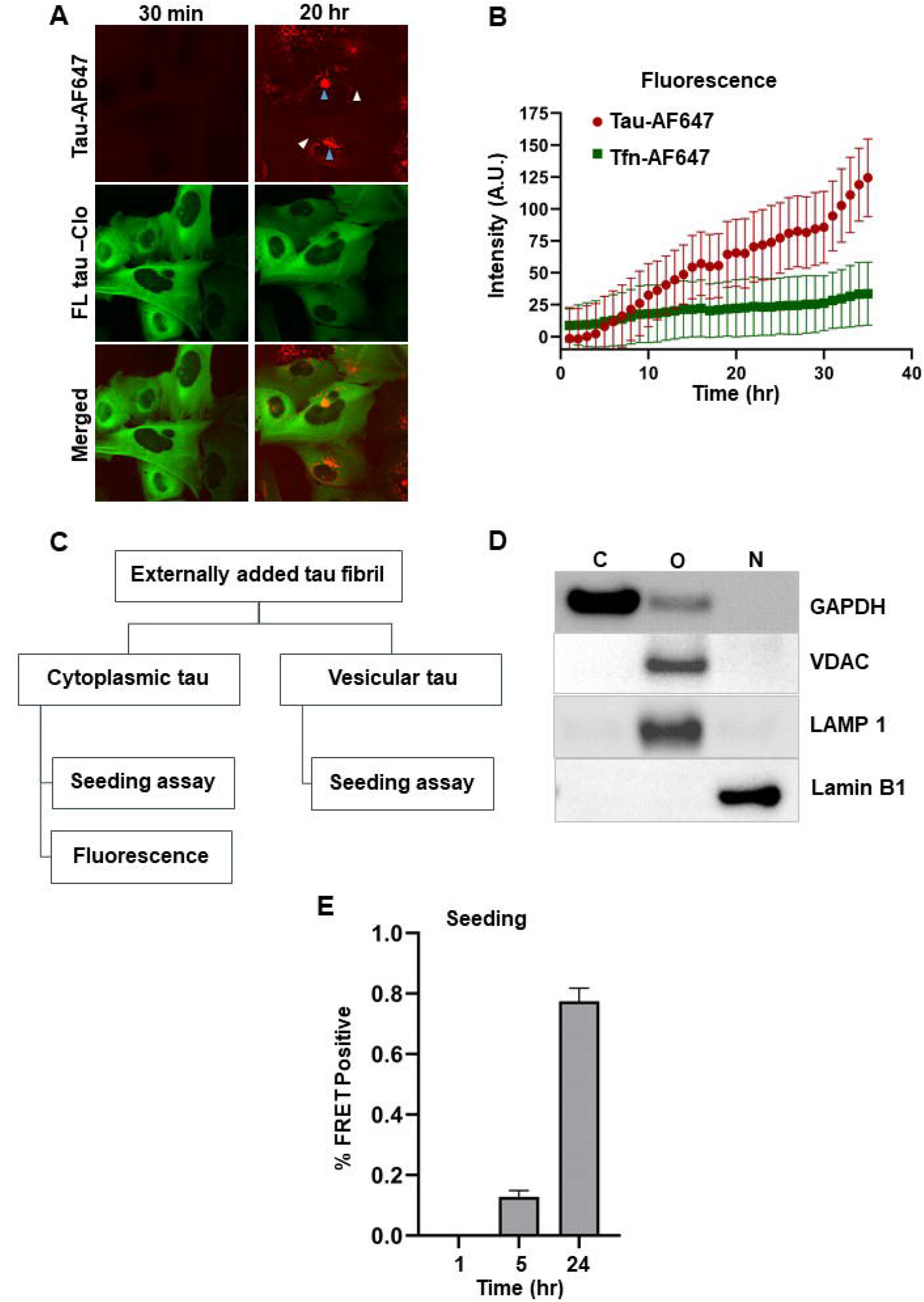
Exogenous tau enters the cytosol. U2OS cells expressing FL tau-Clo (green) were exposed to tau fibrils labeled with AF647 (red), and fluorescence was tracked over time using live cell imaging. (A) Representative images illustrate the increase in the cytosolic AF647 fluorescence between 30min and 20h. Arrowheads indicate cytosolic (white) vs. vesicular (blue) tau. Quantification was based on imaging n>500 cells. (B) Tau-AF647 signal in the cytosol was compared to transferrin (Tfn)-AF647 added to cells. Tau signal increased over time, whereas Tfn signal did not. Fluorescence is indicated in arbitrary units (A.U.). Error bars = S.D. (C) Diagram indicating steps used to quantify exogenous tau seeding in different fractions. Immunoprecipitation was used to measure tau seeding in vesicle fractions, as crude lysate was toxic. (D) Fractions were analyzed by western blot to assess fractionation fidelity. Antibodies against the indicated proteins were used to probe different cell fractions. C: cytosolic; O: organelle; N: nuclear. (E) Cytosol tau seeding was measured using the v2L biosensor. Representative graph shows increase in cytosolic seed over time. The background seeding value from negative control is subtracted from each data point. Seeding assay was performed on 3 independent biological replicates, and each individual measurement was carried out in triplicate. Error bars indicate S.E.M.

### Tau fibrils do not damage vesicles

Prior studies have proposed that tau mediated damage to vesicles might allow leakage into the cytoplasm to initiate seeding (6, 14). To test this hypothesis, we expressed the galectin-3 β-galactoside binding protein Gal3, fused to mRuby3 (Gal3-Rub) to observe the relationship of vesicle rupture to tau seeding. β-galactosides localize to the outer leaflet of the cell membrane and are present in the lumen of endocytic vesicles. Gal3-Rub expressed intracellularly normally is diffusely distributed. However, if an endosome is damaged, Gal3-Rub binds β-galactosides, creating puncta (15)(16). We stably expressed Gal3-Rub in U2OS cells and tracked puncta formation using high content microscopy. Gal3-Rub diffusely distributed in most cells (Fig. 5A). Following exposure of cells to FL tau fibrils tagged with Alexa647 we observed tau in every cell. However, we observed no change in Gal3-Rub puncta formation (Fig. 5A). This was confirmed by quantitation of hundreds of cells (Fig. 5B). We contrasted this with LLOMe treatment to disrupt vesicles, which strongly induced Gal3-Rub puncta formation (Fig. 5B). In summary, we observed no detectable change in overall vesicle permeability upon tau exposure, suggesting that tau fibrils do not significantly damage endocytic vesicles.

**Figure 5:**
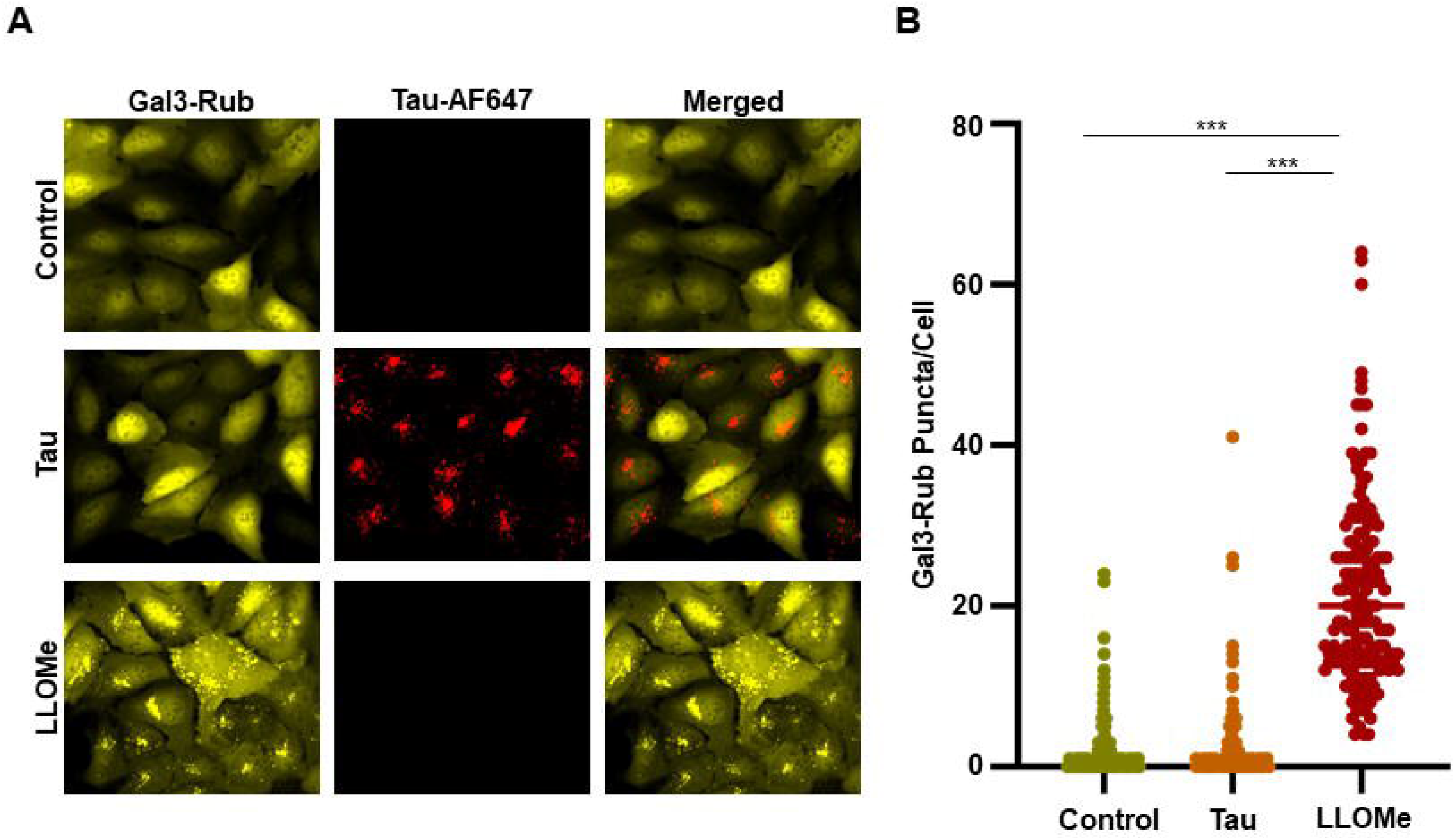
Tau fibrils do not damage vesicles. U2OS cells were stably transfected to express Gal3-Rub as a marker of loss of vesicle integrity. (A) Images of untreated cells (control); cells treated with FL tau-AF647 fibrils; cells treated with LLOMe. No obvious effect of fibril treatment on Gal3-Rub puncta formation was observed, whereas LLOMe produced many positive puncta. (B) Hundreds of cells were imaged after fixation at 24h using high content microscopy, and image analysis was used to quantify the frequency of Gal3-Rub puncta. There was no difference between control cells and those treated with tau fibrils, whereas LLOMe strongly induced Gal3-Rub puncta formation. Data encompasses n∼160-200 cells for each condition. p<.001 for effect of LLOMe.

### Vesicle lysis slightly increases tau seeding

We used live cell imaging to track seeding into cells using expression of FL tau-Clo as a biosensor. We observed seeding into cells with and without coincident Gal3-Rub positive puncta (Fig. 6A,B). Seeding efficiency onto full-length tau is relatively low, so to more easily quantify Gal3-Rub puncta in relation to seeding events, we used the tau RD-clo biosensor. This enabled recording of hundreds of seeding events. We began by observing in an endpoint assay that ∼11% of untreated cells exhibited Gal3-Rub puncta (Fig. 6C). By contrast, ∼35% of cells with seeding exhibited coincident Gal3-Rub puncta, whereas ∼65% did not (Fig. 6C). The higher association of Gal3-Rub puncta with seeded cells led us to test the relationship more rigorously. We used dynamic imaging of cells in culture to identify those that developed tau inclusions after exogenous seeding. We then tracked the cells backwards ∼12h prior to the appearance of tau inclusions to determine the percentage that showed no Gal3-Rub puncta at any time (82%), transient Gal3-Rub puncta (2%), or pre-existing puncta (16%) (Fig. 6D). The large majority of seeded cells did not show any prior evidence of vesicle rupture. The number with pre-existing evidence of vesicle instability (18% total) was slightly higher than the total we observed in untreated cells (12%). To directly compare the seeding efficiency after direct rupture of vesicles, we tested the effect of vesicle rupture by LLOMe vs. seed transduction via Lipofectamine™ 2000. We observed a modest increase in seeding after LLOMe treatment (from ∼1% to ∼5%), however Lipofectamine treatment, which presumably delivers aggregates directly to the cytoplasm, increased seeding to ∼32% (Fig. 6E). We conclude that while vesicle integrity may influence the frequency of seeding events, this is not the primary determinant.

**Figure 6:**
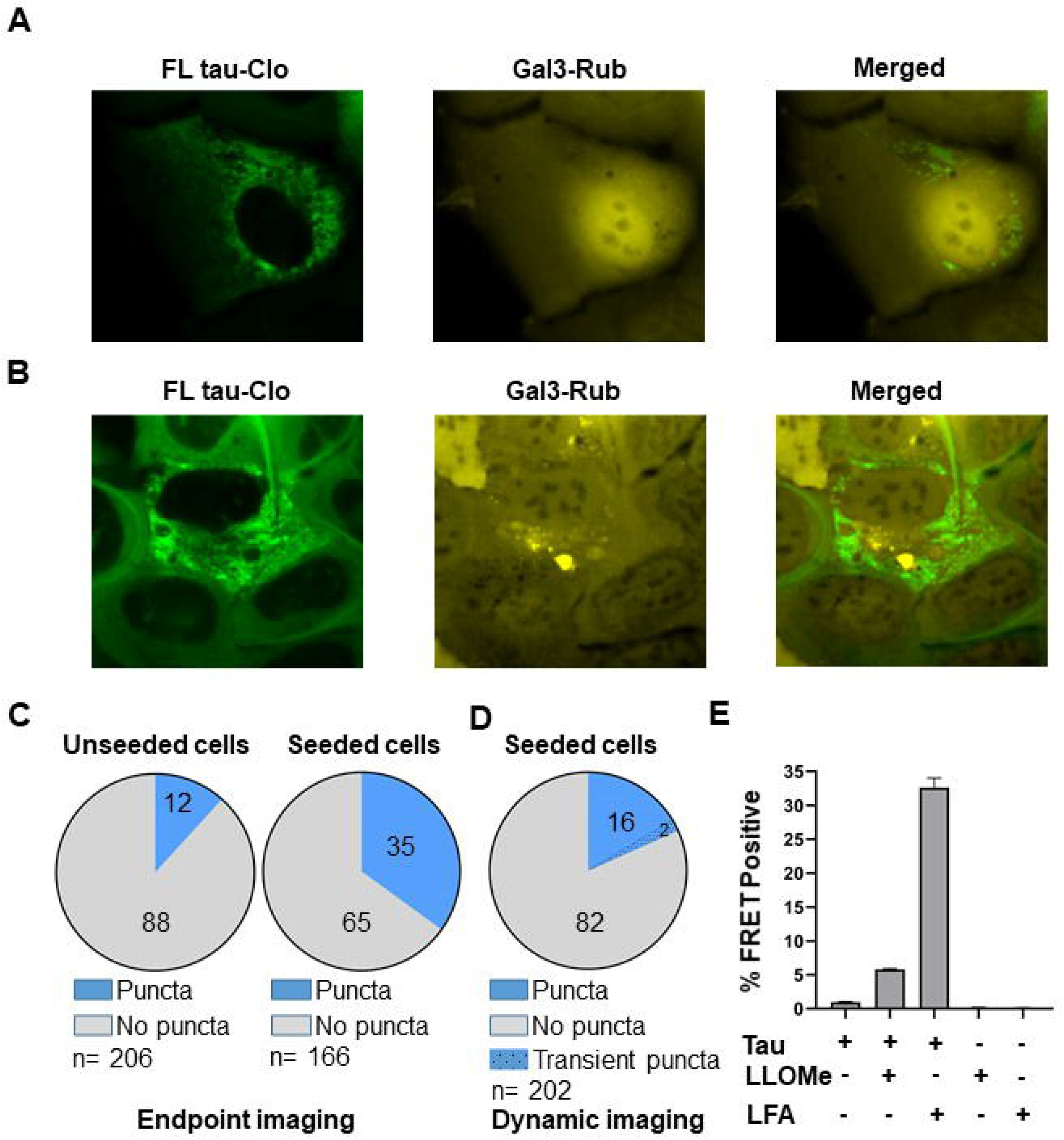
Gal3 positive cells are partially associated with increased seeding. U2OS cells expressing Gal3-Rub were exposed to FL tau fibrils and imaged either in endpoint assays (with cells expressing FL tau-Clo), or via live cell imaging (with cells expressing RD-Clo). (A,B) Representative image showing seeded cells expressing FL tau-Clo (green) without (A) or with (B) coincident Gal3-Rub puncta (yellow). There was no obvious colocalization of tau and Gal3-Rub puncta. (C) Endpoint assay of cells expressing FL tau-Clo with or without exposure to tau fibrils. First chart shows ∼12% of Gal3-Rub puncta in unseeded cells (n∼206 cells). Second chart shows that in cells with evident FL tau-Clo puncta, the incidence of Gal3-Rub puncta increased to ∼35% (n=166 cells). (D) Dynamic imaging of cells expressing tau RD-Clo and Gal3-Rub exposed to fibrils, in which cells exhibiting inclusions were tracked backwards in time to determine the presence or absence of Gal3-Rub puncta. ∼82% of seeded cells showed no Gal3-Rub puncta, whereas ∼18% of seeded cells showed pre-existing (16%) or transient (2%) Gal3-Rub puncta. (E) Comparison of seeding between unmodified application of fibrils, LLOMe, or Lipofectamine 2000 (LFA) -mediated fibril exposure. LLOMe modestly increased seeding, whereas LFA strongly increased seeding. Error bars=S.E.M. Each data point represents a sample analyzed in technical triplicate.

### Distinct mechanisms of tau seed clearance in vesicles vs. cytosol

Our observations were consistent with escape of tau seeds from the vesicular compartment to the cytosol, where seeding occurs. The persistence of tau in each compartment would thus determine the relative efficiency of seeding. Consequently, we evaluated the kinetics of tau seed clearance using purification of seeds from each fraction, coupled with detection using standard v2L biosensor cells (13). We exposed U2OS cells to recombinant FL tau fibrils, followed by fractionation of cells into organelle (vesicle) vs. cytosol fractions. We used immunoprecipitation to test for seeding within the organelle fraction (to avoid toxicity on biosensor cells), and directly assayed the cytosol fraction. In the organelle fraction tau had virtually disappeared by 12h (Fig. 7A). In the cytosol fraction clearance was slightly slower, with tau seeding detectable even at 48h (Fig. 7B). We attempted to use ELISA and mass spectrometry to monitor tau clearance directly, but levels were too low for accurate measurement without using inordinately large amounts of recombinant tau in cultured cells.

**Figure 7:**
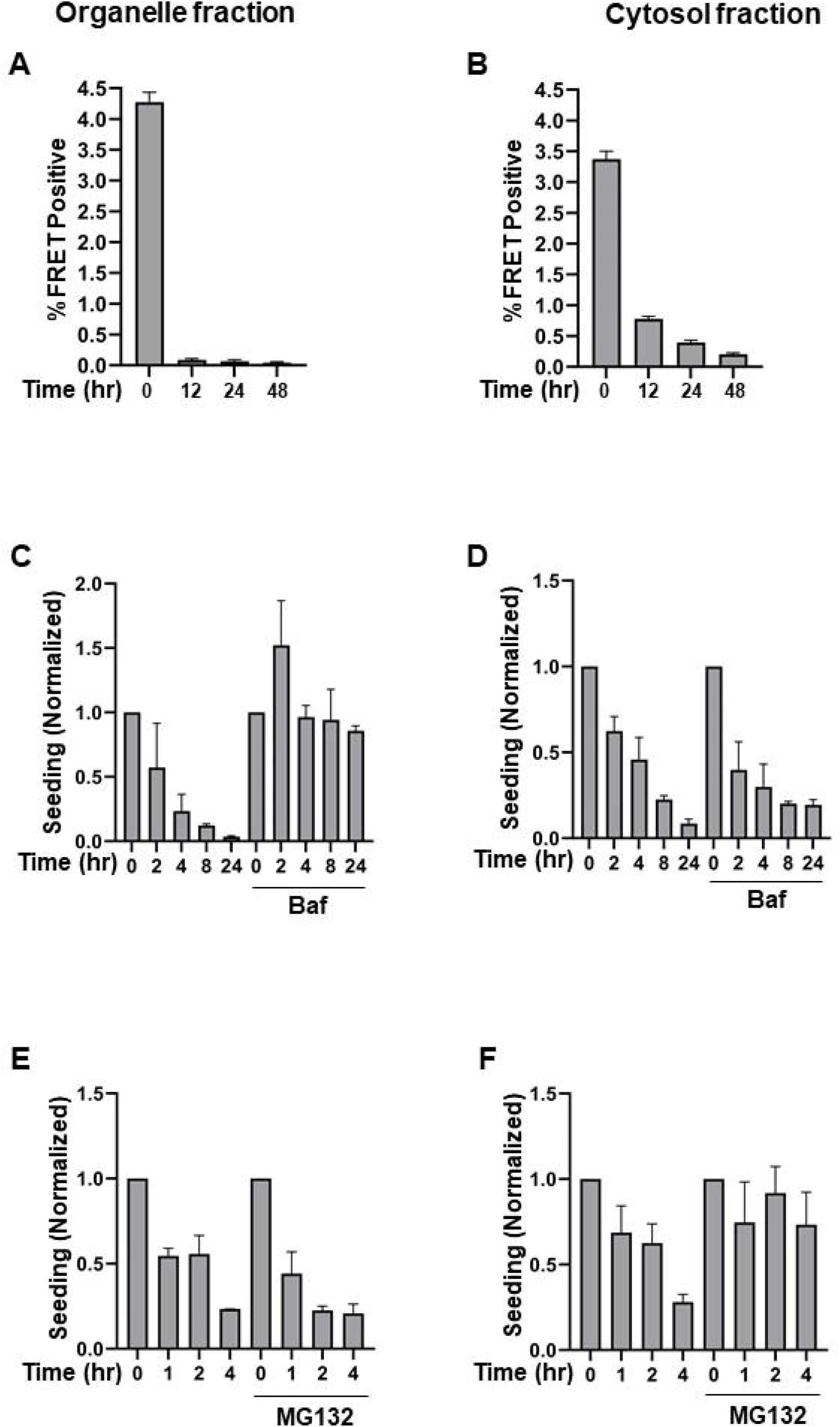
Differential degradation of tau in cytosol vs. vesicles. U2OS cells were exposed to tau fibrils overnight, then treated with trypsin, washed, and incubated for indicated amounts of time. Cells were then subjected to biochemical fractionation to isolate tau in the cytosol vs. organelle (vesicle) fractions. For organelle fractions, seeds were extracted by immunoprecipitation to avoid cytotoxicity of vesicle contents. (A,B) Time course of tau seeding activity present in cytosol (A) vs. organelle (B) fractions reveals degradation in both. (C,D) Bafilomycin (Baf 200nM) was added to block lysosomal acidification. This slowed loss of tau seeding activity in the organelle fraction but had no effect on seeding within the cytosol. (E,F) MG132 (10μM) was added to block proteasome activity. (E) MG132 had no effect on seeding in the organelle fraction, (F) but slowed cytoplasm tau degradation. Assays were performed in technical triplicate. Data is representative of 3 similar experiments for MG132 and Bafilomycin treatment for cytosolic fractions and 2 replicates for organelle fractions. Error bars=S.E.M.

We next tested the effect of inhibitors of the lysosome (bafilomycin) and proteasome (MG132) on seed clearance over 4h, a maximum time point picked to minimize secondary effects. Bafilomycin halted degradation in the organelle fraction (Fig. 7C) but had no effect on the cytosol fraction (Fig. 7D). By contrast MG132 had no effect on organelle clearance (Fig. 7E) but prevented clearance of tau seeds from the cytosol (Fig. 7F). We observed a rapid seed clearance in the organelle fraction. In the cytosol there appeared to be a rapid clearance with t_1/2_ of ∼4h, and a slower phase of 12-24h. Taken together, the data suggest two paths of seed clearance for tau: in the vesicle seeds are degraded via the lysosome, whereas in the cytoplasm seeds are cleared rapidly via the proteasome.

## Discussion

It is unknown how a tau assembly propagates a unique structure from the outside to the inside of a cell. This study has investigated the trafficking kinetics of tau seeds into the cytosol, and their mechanisms of degradation. For ease of labeling and tracking we used recombinant, heparin-induced FL tau fibrils, which undoubtedly lack the same seed conformation that occurs in AD, or post-translational modifications. Second, rather than primary neurons, which might more accurately reflect disease processes, we used an immortalized cell line, U2OS, because it is highly adherent, has a large cytoplasm, and is useful for live cell imaging over days. We observed that most tau taken up by the cell traffics to the endolysosomal system where it is degraded fairly rapidly. A small percentage of assemblies enter the cytosol to seed intracellular aggregation, which appeared often to occur simultaneously at multiple sites. Seeding increased after experimental rupture of endosomes, however this did not appear to be the predominant mechanism, and tau assemblies did not appreciably disrupt vesicles. Most assemblies appeared to be degraded via bafilomycin-sensitive mechanisms in the lysosome, whereas cytosolic seeds were degraded primarily by the proteasome. In summary, our data indicate two major routes into the cell for a tau seed: one into the lysosomal degradation pathway, and the other to the cytosol where recruitment of native tau and template-based replication occurs (Fig. 8).

**Figure 8:**
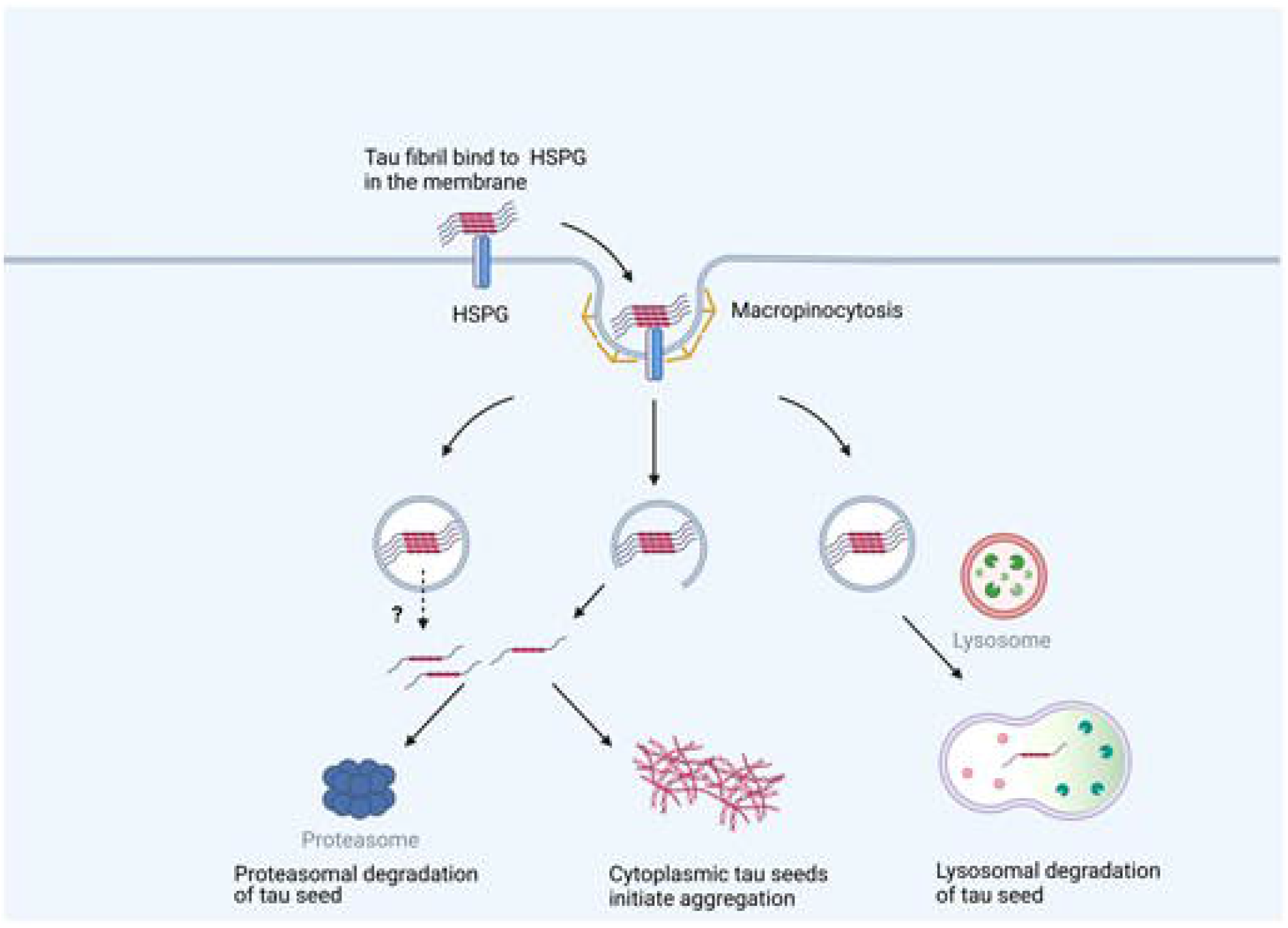
Model of tau trafficking. External tau fibrils bind to the HSPG cell surface receptors which leads to their internalization via macropinocytosis. Most tau within vesicles is directed towards lysosomal degradation. Some tau in this pathway escapes degradation and instead enters the cytosol. This could potentially occur by vesicle rupture, but most tau seeds appear to enter the cytosol in the absence of this process through an unknown mechanism. Once the fibril enters the cytosol, it serves as a template for replication with endogenous tau to amplify the aggregate. Proteasome-mediated degradation in the cytosol accounts for clearance of most tau, although longer-term clearance mechanisms could involve other mechanisms.

### Cytoplasmic tau seeding

In cultured cells, despite widespread tau uptake by macropinocytosis, seeding is relatively inefficient. To overcome this problem in experimental systems, we (12) and others (17) have used lipid-based transduction reagents to improve cytoplasmic delivery of tau assemblies. In biosensor cells expressing FL tau-Clo we observed a seeding efficiency of roughly 1% of exposed cells, despite ∼100% of the cells taking up labeled tau assemblies into the endolysosomal system. We expressed FL tau-Clo constructs that bind the cytoplasmic tubulin network. Within hours after tau seed exposure, however, we observed simultaneous evolution of tau puncta throughout the cytoplasm. The simultaneous appearance of puncta within the cell suggests that there may be a regulatory pathway controlling this process. In data not presented, we tested for a role for the cell cycle, but did not observe any. We have concluded that seeds not visible by microscopy escaped the endolysosomal compartment to recruit endogenous FL tau-Clo away from the tubulin network. Taken together our results imply that following macropinocytosis, only a small subset of tau assemblies wind up in the cytosol where they serve as templates for amplification.

### Endolysosome rupture only partially explains tau seeding

Prior studies have suggested that tau enters the cytosol by rupturing vesicles (6,14). However, using formation of Gal3-Rub puncta as a marker of vesicle integrity, we found no evidence that tau aggregates directly induce rupture. Neither did we observe FL tau-Clo puncta formation in close proximity to Gal3-stained vesicles. Finally, based on live cell imaging of hundreds of cells, in the majority of cases (∼65-80%) we observed no evidence of vesicle rupture in association with seeding events. However, when we specifically tested the relationship of Gal3-Rub puncta to subsequent cell seeding, we observed a slight increase in the percentage of cells that exhibited pre-existing evidence of Gal3-Rub binding (∼35%), relative to cells that did not have evidence of seeding (∼12% of the total overall). This suggested that pre-existing vesicle instability might slightly predispose towards tau seeding. This is consistent with evidence that genes associated with the endolysosomal system are risk factors for AD (18). However, when we treated cells with LLOMe to rupture lysosomes, we only saw a ∼5x increase in seeding, from 1% to 5%, in contrast treatment with Lipofectamine™ 2000 increased seeding ∼35x. This implies that the route by which tau enters the cytosol may impact its seeding efficiency. The contrast of our findings to those of others (6, 14) may reflect our use of different system and purified recombinant tau fibrils. Importantly, a recent study using HEK293 cells and primary neurons also concluded that tau doesn’t damage vesicles (19). We are continuing to investigate the molecular mechanism by which assemblies might cross the vesicle membrane in the absence of rupture, which is the predominant type of seeding event.

### Cytosolic tau seeds are degraded by the proteasome

Our data suggest trafficking of tau seeds into two compartments: the endolysosome and the cytosol. The relative amount of tau protein in both compartments was too low to measure via western blot, ELISA, or even mass spectrometry, but we readily quantified tau seeding activity using biosensor cells. We determined that tau seeds within the vesicle compartment are rapidly cleared. This was blocked by bafilomycin, consistent with lysosomal degradation. By contrast, seeds within the cytosol appeared to have two phases of degradation, a rapid and a slow phase. The rapid phase, with a half-time of ∼2-4h, was blocked by proteasome inhibition with MG132. The persistence of seeds in the cytosol for up to 48h after cell exposure suggests that a small subset may be protected from degradation, with a longer half-life. The role of autophagy in aggregate degradation has been proposed by many studies (20) (21). We do not find evidence consistent with this mechanism for the rapid degradation of most seeds in the cytosol, but it could conceivably play a role in the slower clearance. Since the proteasome must digest single, unfolded proteins, this suggests a role for disassembly factors in this process. One candidate is valosin containing protein (p97/VCP), which has been directly implicated in dominantly inherited tauopathy (22).

### Tau traffics to distinct pools

Our data indicate that two populations of intracellular tau result from macropinocytosis. Most tau enters the endolysomal system for rapid degradation, resulting in relatively inefficient seeding overall. We observed a steady increase of tau in the cytosol over time both by imaging and fractionation studies. Cytosol seeding thus appears to be derived from tau leakage out of the endolysosomal system, whether by vesicle rupture or other mechanisms such as membrane translocation (19, 23). Our conclusions that there exist two paths of tau into the cell, with one leading to intracellular seed amplification, may prove useful in the elucidation of the molecular mechanisms of tauopathy.

### Experimental Procedures

#### Tau fibrillization, labeling and imaging

Recombinant full-length (2N4R) wild-type tau was purified and fibrillized as described previously (3). To label tau, 8μM purified tau fibrils were incubated with .025mg Alexa Fluor 647 (AF647) succinimidyl ester dye (Invitrogen) for 1h at room temperature, and quenched with 100mM glycine for 1h at room temperature. The solution was dialyzed overnight into PBS using dialysis cassettes (Thermo Scientific) to remove unbound dye. Labeled fibrils were stored at 4°C for short term (days to weeks) or at −80°C for the longer term. For live cell imaging assays, labeled fibrils were added to the U2OS cells, and imaged using an IN Cell Analyzer 6000 (GE).

#### Measuring non-vesicular cytoplasmic tau signal

To segment cells within an image, we calculated the image gradient of the FITC channel using the Sobel method, and normalized this output by the gradient magnitude. We then subtracted the resulting normalized image gradient from the original image to enhance cell edges. To remove the remaining image noise, we applied a Gaussian filter with sigma value of 5. Last we calculated the Rosin Threshold from the filtered image and used this threshold value to segment cell boundaries. To segment vesicle boundaries from the Cy5 channel, we applied a lower and upper intensity threshold. For the lower, we calculated the Rosin threshold and ignored all pixels below this value. For the upper (to avoid artifacts from dying cells), we calculated the top 1% intensity and ignored pixels above this value. To compare tau signal inside and outside of vesicles, we calculated tau signal from the Cy5 channel from inside cell and inside vesicle boundaries, or signal inside the cell but outside vesicle boundaries. We then calculated the median intensity for all pixels within these regions. After calculating the median intensity outside of vesicles from the first time point, we background subtracted this value from all subsequent time points.

#### Seeding assay

V2L Biosensor cells (13) expressing tau repeat domain (RD) containing a disease-associated mutation (P301S) fused to mCerulean3 (Cer) or mClover3 (Clo) (tau RD-Clo/Cer) were plated at a density of 10,000 cells/well in a 96-well plate. Recombinant tau fibrils were sonicated for 30s at a setting of 65 (Qsonica Sonicator) and applied to cells at 50μL per well. Cells were then incubated 48h. Tau (80nM) was added directly to the cells after sonication (for naked seeding). Alternatively, Lipofectamine™ 2000 (Thermo Fisher Scientific, Waltham, MA, USA) was used to transduce tau (5nM). After 48h, cells were harvested with 0.05% trypsin, fixed in 2% paraformaldehyde for 10min, and then resuspended in flow cytometry buffer (HBSS plus 1% FBS and 1mM EDTA). We quantified FRET as described previously using an LSRFortessa flow cytometer (7). For each data set three technical replicates were used. For each experiment, a minimum of ∼5000 single cells per replicate were analyzed. Data analysis was performed using FLOWJO version 10 software and GRAPHPAD PRISM version 8.

#### Cell culture and treatments

Human U2OS (ATCC) and HEK 293T cells were cultured in McCoy’s 5A and DMEM medium (Gibco) supplemented with GlutaMAX and 10% fetal bovine serum. Cells were incubated in humidified air with 5% CO_2_ at 37°C and were sub-cultured every 3-4 days. For vesicle permeabilization 1mM LLOMe (Cayman Chemical) was added to the media for 6hrs. Proteasome inhibitor MG132 (Sigma) dissolved in DMSO was used at 10μM final concentration, and the autophagy inhibitor bafilomycin (Sigma-Aldrich) dissolved in DMSO was used at 200nM final concentration.

#### Western blot

Lysates were mixed with 4X SDS buffer and run on a NuPAGE 10% Bis-Tris Gel at 100V. The gel was then transferred onto Immobilon P membrane for 1hr at 20V using a semi-dry transfer apparatus (Bio-Rad). The membrane was blocked with 5% Blotto (Bio-Rad) in TBST for 1.5 hour before primary (GAPDH:1:5000, Lamin B1:1:1000, Lamp 1 – 1:1000, VDAC:1:500) antibody was added at specific dilutions and incubated on a shaker overnight at 4°C. The membrane was then washed three times with TBST at 10min intervals. It was re-probed with (goat anti-rabbit/mice) secondary antibody for 1.5hr at room temperature. The membrane was washed four times with TBST and exposed to ECL Prime western blot detection kit (GE Lifesciences) for 2 min. Blots were imaged with a Syngene digital imager.

#### Cellular fractionation

Fractionation was guided by published protocols (8)(9). Cells were trypsinized and resuspended in 5mL of culture medium. They were centrifuged at 500 x g for 10 min at 4°C, the supernatant was discarded, and the pellet washed with 500μL of ice-cold PBS. ∼250μl (depending on pellet size) of ice-cold lysis buffer (150mM NaCl, 50mM HEPES (pH 7.4), 25 μg/mL digitonin,1M hexylene glycol) was added to the pellet and then incubated on end-over-end rotator for 10-15min at 4°C. The sample was centrifuged at 2000 x g for 10min at 4°C. The supernatant was collected, and the pellet processed for the next step. The supernatant was clarified by centrifugation at 18000 x g for 20min. Supernatant was collected as the cytosol fraction.

The pellet was washed with wash buffer (150mM NaCl, 50mM HEPES (pH 7.4)) and then 250μl of ice-cold buffer (150mM NaCl, 50mM HEPES (pH 7.4), 1% IGEPAL (v:v), 1M hexylene glycol) and resuspended by vortexing. It was incubated on ice for 30min, then centrifuged at 7000 x g for 10min at 4°C. The supernatant was collected. This fraction contains the proteins from all membrane-bound organelles (endosomes, mitochondria, endoplasmic reticulum, Golgi, etc.) except nuclei. The pellet was washed with wash buffer and then 250μl of ice-cold buffer (150mM NaCl, 50mM HEPES (pH 7.4), 0.5% sodium deoxycholate (w:v), 0.1% sodium dodecyl sulfate (w:v), 1M hexylene glycol, and 7μL of Benzonase (25000 units/ml) was added. It was incubated on an end-over-end rotator for 30min at 4°C to allow complete solubilization of nuclei and digestion of genomic DNA. It was then centrifuged at 7800 x g for 10min at 4°C. The supernatant was collected. This fraction contains the nuclear proteins. 1X Roche EDTA-free complete protease inhibitor was added to all buffers.

#### Seed degradation assay

The U2OS cells were incubated with tau fibrils for 16h. The next day, cells were washed with PBS for 2min, and a .05% trypsin wash (30sec) was used to digest attached tau, followed by quenching with a 3min wash with McCoy’s 5A media. Cells were then incubated for the defined amount of time in media without tau, trypsinized and fractionated to collect the cytosolic or organelle fractions, which were used for seeding assays.

#### Immunoprecipitation

75μl of Dynabeads Protein A (Thermo Fisher) were washed according to the manufacturer’s protocol and incubated with 15μg of tau polyclonal antibody raised against tau RD TauA (13), for 1h at room temperature. The beads were washed with PBST. The beads were added to 75μg of lysate (organelle fraction) and incubated with rotation overnight at 4°C. The next day beads were washed with PBST and protein eluted in low pH elution buffer (Pierce). The reaction was neutralized with 1:10 1M Tris pH 8.5 in a final volume of 120μl.

## Data Availability

All data generated and analyzed during this study are included in this article.

## Supporting information

This article contains supporting information.

## Acknowledgements

This work was supported by the Crowley Foundation, the Cure Alzheimer’s Foundation, the Rainwater Charitable Foundation, the Hamon Foundation, and the NIH (WU-16-376-MOD-5). We also thank the Moody Foundation Flow Cytometry Core Facility and the Sequencing Core Facility at UT Southwestern.

## Author contributions

MID and SK designed research; SK, ARV, DAD, VAP, OMK, CLW performed research and/or contributed reagents; MID and SK wrote the manuscript.

## Funding and additional information

This work was supported by the Crowley Foundation, the Cure Alzheimer’s Foundation, the Rainwater Charitable Foundation, the Hamon Foundation, and the NIH (WU-16-376-MOD-5).

## Conflict of interest

The authors declare no conflict of interest.

## Figure legends

**S1Movie:** Recombinant FL tau fibrils labeled with Alexa Fluor 647 were applied to the U2OS cells expressing FL Tau-Clo (green) and imaged using a GE IN Cell Analyzer 6000 to track tau aggregation formation over time. The images taken at different time points post fibril addition are used to make this movie (same cell is shown in Fig 1D). Rapid and simultaneous development of tau inclusions throughout the cytoplasm is evident.

## Notes

### Competing Interest Statement

The authors have declared no competing interest.

